# Evidence for the key roles of the *Pseudomonas syringae* mobilome in shaping biotic interactions

**DOI:** 10.1101/2024.03.19.585818

**Authors:** D. Holtappels, G.E.J. Rickus, T. Morgan, R. R. de Rezende, B. Koskella, P. Alfenas-Zerbini

## Abstract

The mobilome, defined as the collection of mobile genetic elements within a bacterial genome, plays a critical role in the adaptation of bacteria to abiotic and biotic drivers. In particular, prophages have been reported to contribute to bacterial resistance to virulent bacteriophages, the competitive interaction of bacterial hosts within microbial communities, and in pathogenicity and virulence. It is therefore critical to better understand the role of prophages in distributing genes and functions within and among bacterial species to predict how bacteria adapt to their biotic environment. *Pseudomonas syringae* offers an ideal study system to ask these questions both because of its broad range of lifestyles (spanning from environmental growth to plant pathogens) and its high intraspecies diversity. To examine the role of the mobilome in this species complex, we compared 590 genomes available from public databases and annotated the defense mechanisms, effectors, and prophages in the genomes. We found that this species complex has an elaborate phage pandefensome consisting of 139 defense mechanisms. Host-associated *P. syringae* isolates were found to have both elaborate phage defensomes and effectoromes. Assessing taxonomical signatures of the observed prophages uncovered broad differences in the types and numbers of genes encoded by different phage families, emphasizing how the evolutionary advantages conferred to hosts will depend on the prophage composition and offering insight to how these genes might disperse within a community. Our study highlights the intimate association of phage families with their hosts and uncovers their key role in shaping ecology for this widespread species complex.

**Significance statement:** The bacterial accessory genome, including the mobilome and prophages, plays a critical role in shaping bacterial adaptation to abiotic and biotic drivers. These prophages are widespread across bacterial taxa and likely maintained because of their evolutionary advantage. Our ability to predict how a bacterial population will evolve over time requires a better understanding of where key functional traits arrive. To address this question, we assessed prophage-encoded phage defenses and effector across *Pseudomonas syringae*. We show that prophages carrying these genes belong to specific phage taxa with differences in the types of genes encoded. This emphasizes the evolutionary advantage of these prophages, offering a framework to uncover how these genes disperse within microbial communities and their role in pathogen evolution.

## Introduction

Bacteria are continuously under evolutionary pressures from abiotic and biotic interactions within a given environment (1, 2). To respond to these complex dynamics, bacteria must evolve rapidly. In addition to their often high mutation rates and rapid generation times (especially compared with most eukaryotes), bacteria have a remarkable degree of genome plasticity resulting from the acquisition of new genetic material (e.g. genes encoded on mobile genetic elements such as phages and plasmids) from the environment or members of the community (3, 4). In the case of bacterial pathogens, for example, the acquisition of virulence genes embedded in pathogenicity islands through horizontal gene transfer, including the dispersal of genes through viral vectors, can lead to host range expansion and key changes in virulence (5). Importantly, host-associated bacteria are not only entangled in a coevolutionary arms race with their eukaryotic host, but also with their bacteriophages (6, 7). As lytic phages lyse and kill bacterial cells for successful reproduction, they impose strong evolutionary pressure on susceptible bacterial populations, resulting in diverse resistance strategies that continually reshape who is infecting whom (8). Recent advances in our understanding of phage defense mechanisms make clear that horizontal gene transfer can drive distribution of these mechanisms (9, 10). As such, we can hypothesize that the mobilome, defined as all types of mobile genetic elements within a bacterial genome, is a critical driver of bacterial adaptation, not only in its interactions with the host, but also its resistance to viral predation (2).

As part of the mobilome, prophages and viral satellites, among other types of genomic parasites, can act as carriers for the dispersal of genetic information between bacteria in a community. Often these genes offer the host an evolutionary advantage, referred to as lysogenic conversion (11), as these genes are involved in metabolic pathways, influence the outcome of biotic interactions within the community (12, 13), or shape the bacterium-host interaction (11). For example, several prophages are described that carry auxiliary metabolic genes, such as those involved in the carbohydrate metabolism and two-component transporter for the uptake of specific compounds (14), and direct microbial interactions can be modulated by prophage-encoded antimicrobial resistance genes (15). These genomic parasites not only modulate bacterium-bacterium interactions, but also shape the outcome of phage-bacterium interactions themselves. As prophages are vulnerable to other superinfecting phages, they have evolved to carry mechanisms that aid in surviving phage predation, referred to as superinfection exclusion mechanisms (as reviewed elsewhere (16)). Alongside these specialized mechanisms, prophages can also encode restriction enzymes (17, 18) and CRISPR Cas phage defense mechanisms (19). Prophages can also carry resistance mechanisms that act beyond the individual cell to protect bacteria at the population level, such as abortive infection mechanisms (Abi). A recent example of such an Abi encoded by a prophage is the BstA mechanism that effectively offers a population-wide protection against phage predation, including an anti-BstA hampering the activity of BstA that allows the prophage to go into a lytic life cycle (20). Lastly, prophages are capable of drastically shaping bacterium-host interactions through incorporation of genes involved in the pathogenesis of the host, or modulation of the bacterium-host interaction. For example, *Vibrio cholerae*, depends on the presence of prophages encoding the cholera toxin (CTX) for their pathogenicity (21). Further prophage-encoded proteins underlying pathogenesis include the shiga toxin in Shiga toxin-producing *Escherichia coli* (STEC), the diphtheria toxin in *Corynebacterium diphtheria,* the botulinum toxin in *Clostridium botulinum,* and the SopE effector in *Salmonella enterica* (22). In *Staphylococcus aureus,* beta-haemolysin-converting prophages even encode genetic clusters that aid the bacterial host to evade the immune system of the eukaryotic host and are associated with disease severity in chronic inflammatory diseases like chronic rhinosinusitis (22, 23). These examples illustrate the intricate interaction between bacteria and their genomic parasites and how they shape the adaptability of their host, ranging from the abiotic to biotic interactions. further illustrating the intricate interaction between bacteria and their phages.

Many open questions remain about the distribution and phylogenetic conservation of mobile genetic elements encoding bacterial virulence genes and phage defense mechanisms. One promising avenue to address these questions is using comparative genomics among highly related species that differ in both their biotic interactions and prophage content to infer key prophage-encoded functions. *Pseudomonas syringae* is a highly diverse species complex that consists of thirteen different phylogroups and hence is characterized by a high interspecies diversity (24). This species includes both highly specialized phytopathogens causing disease in a myriad of commercially and environmentally relevant plants, as well as isolates that are non-host associated (25–27). The mobilome of this species complex has been shown to play an intricate role in the adaptation of *P. syringae* to its abiotic and host environment. Plasmids, for example, are described to aid in the resistance against xenobiotics (i.e. copper chemicals and antibiotics such as streptomycin) across different isolates (28), and the type III effector proteins (key drivers of host specialization) are often found to be encoded on pathogenicity islands and plasmids (29–32). As such, these genes are capable of horizontal transfer among unrelated strains and species, highlighting the key role of the mobilome in pathogenicity. Recent insights to the relationship between the pathogenesis of *P. syringae* and its prophages demonstrated that prophages also encode effector proteins. *P. syringae* pv. *morsprunorum* was found to encode HopAR1 effectors embedded in prophage sequences that were found to excise and circularize, leading to successful transferred from one bacterium to another (33). Although data such as these clearly demonstrate the potential importance of prophages in the pathogenesis of *P. syringae,* there has been relatively little work on if and how prophages carry phage defense mechanisms.

To address this critical knowledge gap, we characterize the phage defensome, the effectorome, and prophage-like elements encoded by 590 publically available *P. syringae* genomes within the *P. syringae* species complex, focusing on the distribution and potential interaction of these genetic elements. We specifically evaluated the likeliness that a defense mechanism is distributed through horizontal gene transfer and confirmed our findings through sequence alignments and phylogenetics. We establish a network of prophage-like elements between the different phylogroups and perform correlation analyses between the phage defensome and the prophages encoded by isolates within our dataset to grasp the vastness of mobile genetic elements within the *P. syringae* species complex and the interaction between these drivers.

## Results

### The *P. syringae* species complex represents a diverse panel of isolates with elaborate phage defensomes and effectoromes

We extracted over 680 assemblies of *Pseudomonas syringae* genomes from NCBI, corrected for duplicates and evaluated their completeness by means of a BUSCO analysis. As a result, we narrowed the dataset down to 590 genome assemblies with a BUSCO score greater than 0.95. These genome assemblies were used to calculate the phylogenetic relationship between the different isolates within NCBI. Interestingly, there are significant differences in the assembly size between the different phylogroups (Fig 1 A – Tukey-Kramer test with corrections for multiple comparisons p-value < 0.05). Isolates grouped in phylogroup 1, 6 and 13 have the largest genomes with an average of 6.2 Mbp, 6.1 Mbp and 6.4 Mbp, respectively, followed by phylogroups 2, 5 and 7 with an average of 6.0 Mbp, 5.98 Mbp and 5.93 Mbp, respectively. Members of phylogroup 10 have the smallest genome size with an average of 5.56 Mbp (Figure 1A).

**Figure 1.**
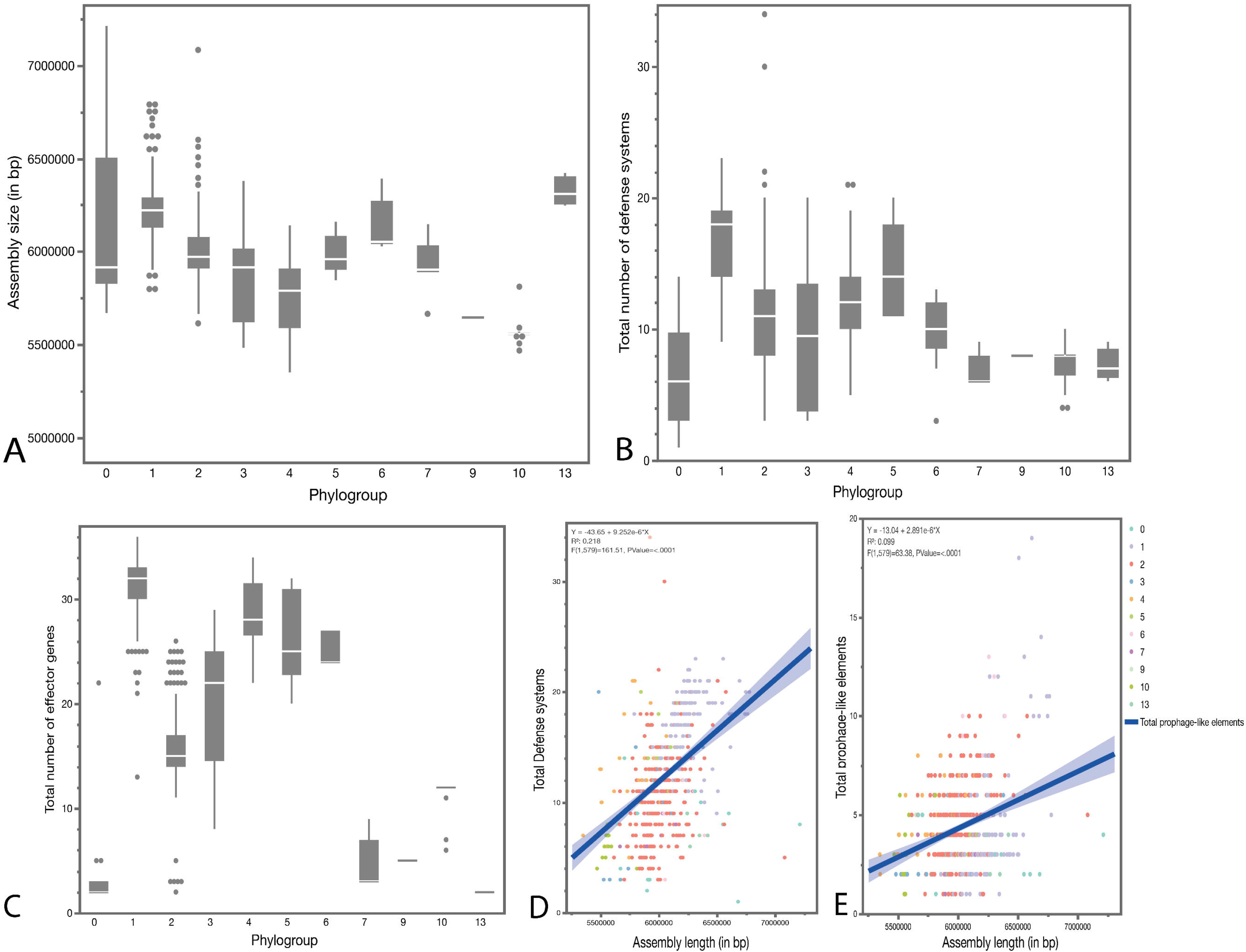
Basic characteristics of the bacterial draft genomes organized by phylogroup as visualized as outlier box plots. A. Total assembly size of the different draft genomes per phylogroup. Phylogroups 1, 6 and 13 had on average the largest genome size (Tukey test with correction for multiple comparison p-value < 0.05) B. Total number of annotated phage defense systems per genome organized by phylogroup. Phylogroup 1 and phylogroup 5 encoded on average the most phage defense mechanisms (Tukey test with correction for multiple comparison p-value < 0.05), and C. Total number of effector genes annotated in the draft genomes per phylogroup with phylogroup 1, 4 and 5 encoding the highest number of effector proteins as previously described by Dillon et al. 2019(21). D. and E. Linear regression analysis of the total number of defense systems and the total number of prophage-like elements in function of the assembly size in bp, respectively. Phylogroups are color coded as follows: PG0 in teal, PG1 in violet, PG2 in orange, PG3 in blue, Pg4 in yellow, PG5 in light green, PG6 in pink, PG7 in purple, PG10 in olive, and PG13 in dark green. The correlation coefficient R^2^ was 0.244 for the defense mechanisms and 0.099 for the effector genes indicating weak correlations between the variables. The p-value (F-test) was <0.0001.

Next, we screened the genomes for over 60 different types and 150 subtypes of defense mechanisms, as currently reported in literature, as well as a panel of the most important effector genes (Figure 1B and Figure 1C). We found that throughout the species complex of *Pseudomonas syringae*, isolates encoded on average 12 defense mechanisms in their genomes and 20 effector genes. The total pandefensome of *P. syringae* consisted of 139 types of defense mechanisms. Based on our annotations, septu mechanisms were the most abundant in the *P. syringae* pandefensome with 911 annotated mechanisms, followed by Gibija (n=546) and Type IV and Type II restriction modification systems (n=529 and n=465, respectively). The least abundant mechanisms were related to other abortive infection mechanisms (Abi), CRISPR-Cas, and BREX (n=1) (Supplementary Fig. 1).

In the case of the effector genes, most of *P. syringae* isolates in the dataset encoded HopB and HopBV (n=581). Other abundant effectors genes were *avrE* (n=555)*, hopAA* (n=549), *hopAH* (n=549), *hopM-shcM* (n=545), *hopAG* (n=520), and *hopAI* (n=507). Rare effector genes were *hopBG* (n=3), *hopU* (n=3), *hopBT* (n=2), and *hopBW* (n=1). Our correlation analysis of the total number of phage defenses and effector genes encoded per genome as a function of the genome size demonstrated that there is a general positive correlation between these variables (Fig1D-E). In this case the correlation between defense mechanisms and genome size was stronger (R^2^ 0.218, p-value <0.01) suggesting that larger genomes carried more defense mechanisms (Figure 1D). Within our model, there was a significant effect of the bacterial phylogroup, but not of the assembly length, nor the interaction term between phylogroup and assembly length.

In terms of the number of defense mechanisms that were encoded by the different phylogroups (Fig 1B), we found that Phylogroup 1 isolates tended to encode the most phage defense mechanisms (average of 17 mechanisms per genome Group A), followed by phylogroups 5 (14 defense mechanisms Group AB) and 4 (12 defense mechanisms Group B; Tukey-Kramer test with correction for multiple comparison, p-value <0.05). Phylogroup 2 (Group BC) and phylogroup 3 (Group BCD) consist of important pathogenic isolates of *P. syringae,* encoding an average of 11 defense mechanisms. Isolates that were not assigned to a phylogroup in literature (Phylogroup 0) encoded on average 6 defense mechanisms per genome (Group D). Overall, our analyses demonstrated that the number of defense mechanisms was shaped by genome size and depended on the phylogroup of the bacterial host (Supplementary Figure 2).

In contrast, when we examined the effector genes, we found a significant interaction between phylogroup and assembly length (genome) size (p-value = 0.0198). Including this interaction term, we could not define a significant difference in the number of effector genes encoded between the different phylogroup. However, within the absence/presence matrix of the different effector genes, we distinguished fourteen different clusters (Fig. 2). Based on this clustering, we observed some degree of conservation in the effectoromes within phylogroups. Despite this conservation, we found that isolates of phylogroup 1 clustered across five different groups according to their effector profiles. Similarly, isolates of phylogroup 2 grouped across six different clusters and isolates of phylogroup 4 and 6 were characterized by quite divergent effector profiles.

**Figure 2.**
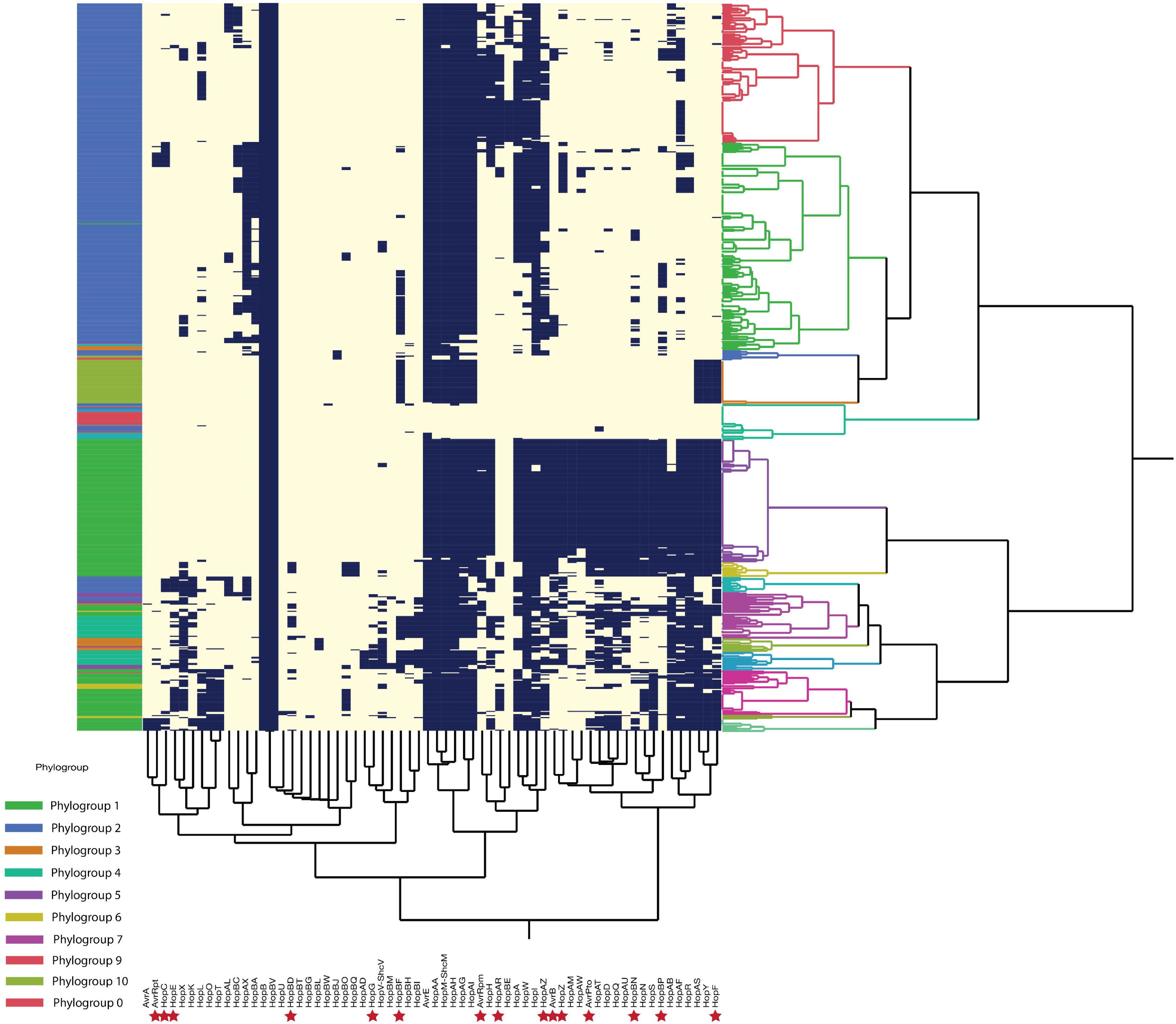
Hierarchical clustering of the absence/presence matrix of the different effector genes (x-axes) annotated in the different draft genomes (y-axes). Phylogroups are indicated on the left-hand panel: PG1 green, PG2 blue, PG3 orange, PG4 teal, PG5 purple, PG6 yellow, PG7 violet, GP9 maroon, PG10 olive, and PG0 in red. The different clusters as determined by the hierarchical clusters are shown on the right-hand. Fourteen clusters were calculated and color coded. Effectors that were annotated in prophage genomes were indicated by a red star.

Moving from the Phylogroup level to the individual genome level, we observed that certain isolates encoded multiple copies of the same class of defense mechanism. We found that there were up to three copies of PfiAT, RM Type I, RosmerTA, and Shedu, and four copies of Gabija, Lamassu, RM Type II, and RM Type IV incorporated in the genomes of *P. syringae* isolates. In the case of Septu, up to five copies were annotated (BS2900, H2a, J6, MAFF 212171, PA-5-7A, PS25, SeraOz_1_1r, and USA007 all isolates within PG1). This potentially demonstrates that *P. syringae* isolates tend to specialize in a specific defense mechanism rather than generalize. To test this hypothesis, we plotted the copy number of a given defense mechanisms as a function of the total number of defense mechanisms encoded per genome. Across the evaluated defense mechanisms, we found in general poor, yet significant, correlations between the copy number of defense mechanisms and the total number of mechanisms encoded per isolate (R^2^ < 0.1, p-value <0.05). Thus, we can assume that the total number of defense mechanisms is a relatively poor, yet significant, predictor for the copy number of these mechanisms, suggesting that the isolates are not specializing in these types of defenses, but rather generalize in a wide diverse panel of mechanisms. However, In case of DarTG, Gabija, Lamassu, Olokun, PD, PfiAT, Restriction modification systems, Septu, Shango, and Wadjet, we calculated relatively higher correlations between their copy numbers and total defenses encoded (R^2^_Gabija_ = 0.31, R^2^_DarTG_ = 0.23, R^2^_Lamassu_ = 0.20, R^2^_Olokun_=0.23, R^2^_PD_=0.15, R^2^_PfiAT_ = 0.29, R^2^_RM_ = 0.50 R^2^_Septu_ = 0.13, R^2^_Shango_ = 0.21, and R^2^_Wadjet_ = 0.21; p-value <0.01).We can thus hypothesize that the more defense mechanisms an isolate encodes, the more of these mechanisms are incorporated into the defensome, especially in the case of restriction modification systems (Supplementary Figure 3). Notably, isolate CFBP13574 from phylogroup 2 was annotated to encode a CRISPR-Cas Class IF mechanism along with only three other defense mechanisms, which is in high contrast to the average of eleven mechanisms for this phylogroup.

### Overall patterns in the *P. syringae* phage defensome are phylogenetically conserved with clear evidence of horizontal gene transfer for certain mechanisms

To investigate the phylogenetic conservation of overall patterns in the phage defensome and of *P. syringae*, we summarized all the annotated defense mechanisms according to the individual isolate and clustered the strains based on the patterns of absence/presence of a specific mechanism (Fig. 3). Then, to evaluate the similarity of the patterns observed in the different defensomes within and between phylogroups based on the absence/presence of a specific mechanism, we performed an Analysis of Similarity (ANOSIM). These analyses uncovered striking differences in the defensomes based on phylogroup (R = 0.274 and p-value = 0.001), suggesting a higher similarity of the defensome within phylogroups than among phylogroups. Despite the phylogenetic relation in the defensome profiles, we observed clear randomness in the profiles within the clusters, hinting towards the horizontal dispersal of certain phage defense systems. To test the likelihood that a specific defense mechanism is acquired horizontally rather than vertically, we calculated the normalized recombination phylogenetic diversity statistic (nRecPD) for each defense mechanisms that is represented at least ten times in the dataset (to ensure statistical rigor). The closer the nRecPD value is to zero, the more likely it is acquired horizontally rather than vertically through phylogeny (34). Supplementary Figure 4 summarizes the individual statistics for each defense mechanism (n>10). Based on these measures, 112 out of 139 mechanisms (total number of mechanisms annotated in the species complex) were found to be likely distributed in the *P. syringae* species complex by means of horizontal gene transfer, highlighting horizontal distribution as a dominant driver of diversity in this pathosystem.

**Figure 3.**
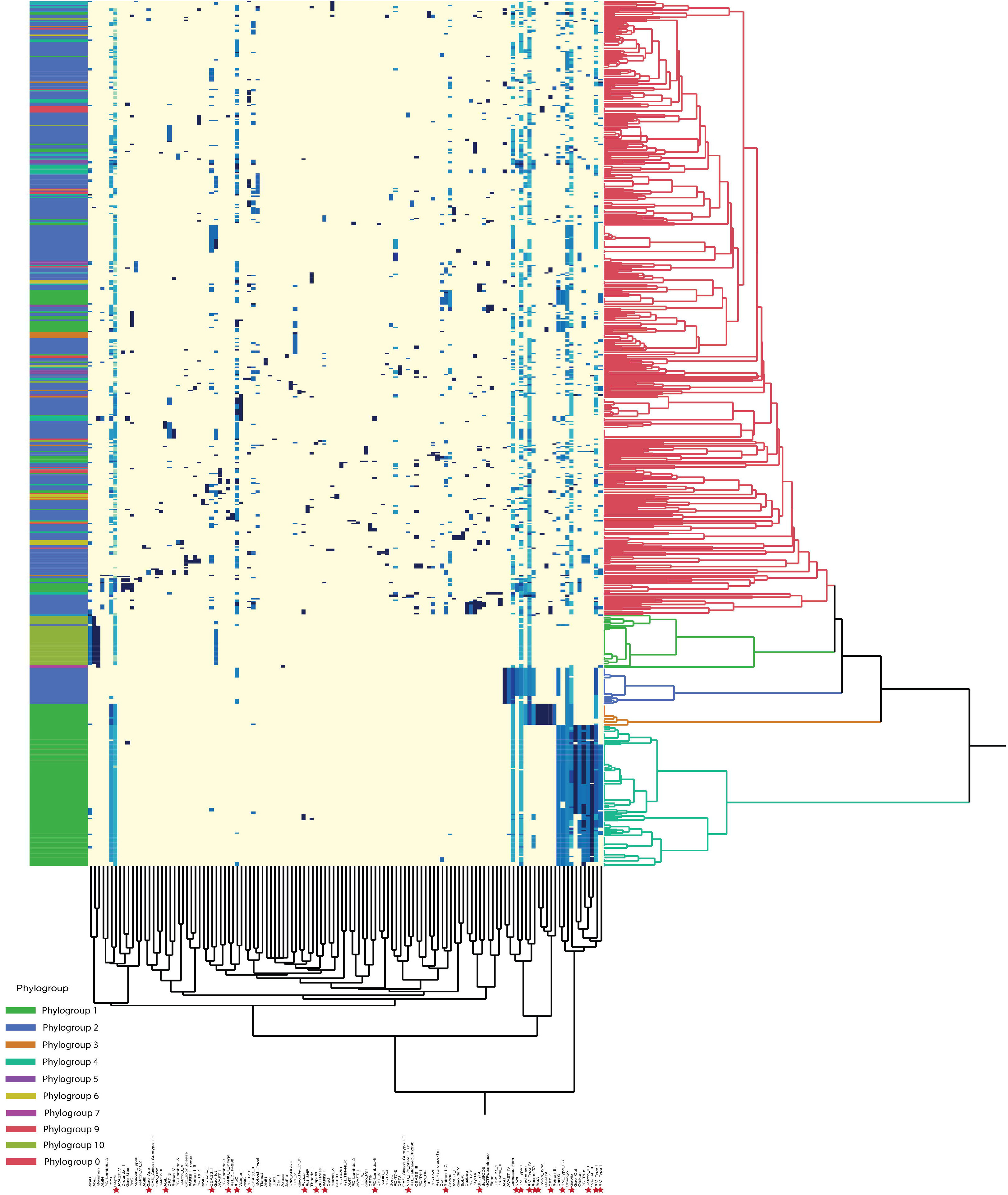
Hierarchical clustering of the absence/presence matrix of the different phage defense systems (x-axes) annotated in the different draft genomes (y-axes). Phylogroups are indicated on the left-hand panel: PG1 green, PG2 blue, PG3 orange, PG4 teal, PG5 purple, PG6 yellow, PG7 violet, GP9 maroon, PG10 olive, and PG0 in red. The different clusters as determined by the hierarchical clusters are shown on the right-hand. Five clusters were calculated and color coded. Defense mechanisms that were annotated in prophage genomes were indicated by a red star.

Because the nRecPD is based on the absence/presence of a mechanism and independent of the actual sequence homology, we built amino acid sequence alignments of the defense mechanisms at subfamily level with an nRecPD value lower than 0.25 and found that there was an overall effect of phylogeny in line with the overall pattern from presence/absence data (Supplementary Figure 5). In general, there is a clear conservation of the different mechanisms according to phylogroup, yet we observed some scattering of the isolates as for most of the defense mechanisms, the same subclass of mechanism was shared between isolates from phylogroup 1 and phylogroup 2 (AVAST II, Druantia II2, Gao Qat 3, Rst, SanaTA, and Wadjet I4). In case of Druantia II2, mechanisms annotated from isolates belonging to phylogroup 3 and 4, as well as 6 and 2 clustered together. Similarly, for Kiwa, phylogroup 5 and 2 as well as 2 and 3 clustered in the same group, respectively. ShosTA, SspBCDE and Wadjet I2 were shared by phylogroup 1, 2 and 4. Notably, when multiple copies of a specific mechanism were encoded, we observed that isolates had acquired different variants of the respective mechanism in the case of Wadjet I4, SspBCDE, SanaTA, Rst, and Olokun, suggesting that isolates diversified in their defense repertoire. Hence, we confirmed our hypothesis at sequence level that these mechanisms are likely to be dispersed through horizontal gene transfer.

### The *P. syringae* species complex carries a wide diversity of genomic parasites

The 590 high-quality bacterial genome assemblies were subjected to a prophage sequence search using PHASTER (35). A total of 2595 prophage-like sequences were detected in nearly all genomes analyzed, except for *P. syringae* pv. *actinidiae* C26 (GCA_002175055.1). Most of the prophage-like sequences were incomplete (1235/2595 = 47.6%), followed by intact (860/2595 = 33.1%) and questionable (500/2595 = 19.3%). Among the *P. syringae* genomes harboring prophage-like sequences, the number of elements ranged from 1 to 19. Only 92 bacterial genomes (92/589 = 15.6%) were free of intact prophages (putatively active phages). Similarly to the defense mechanisms, we plotted the total number of prophage-like elements as a function of genome size and observed a weak positive correlation between the two variables (Figure 4A).

**Figure 4.**
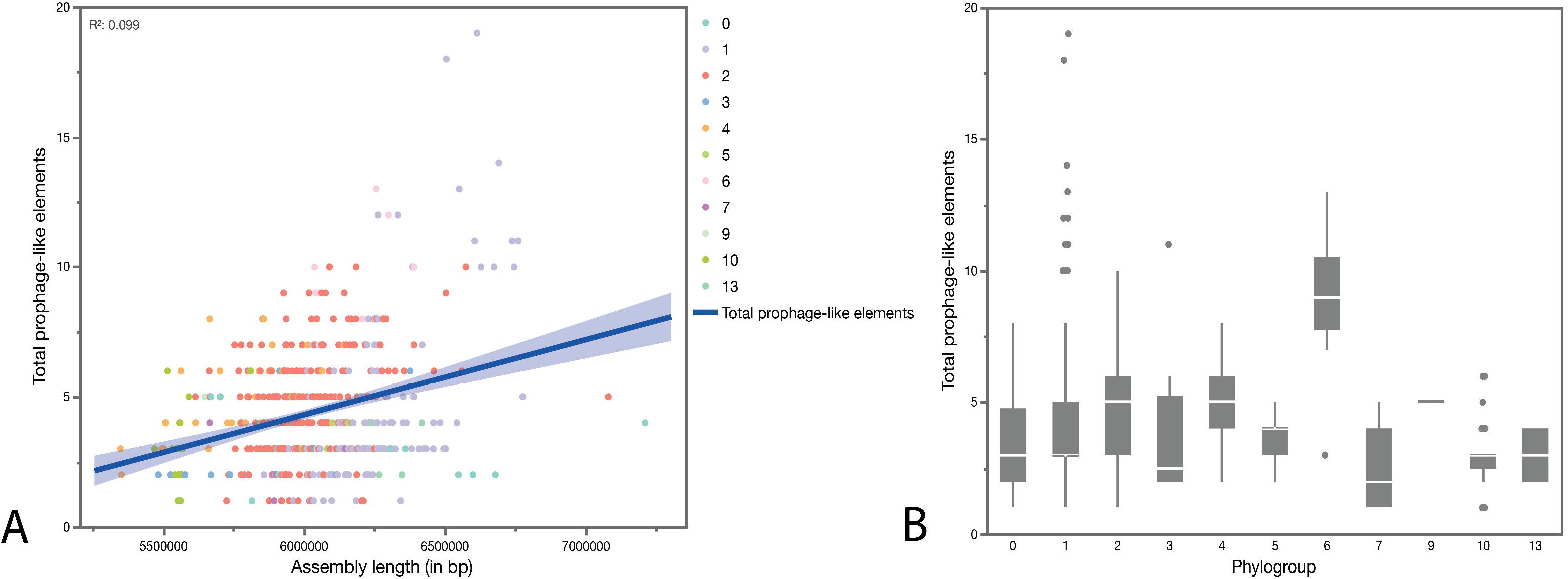
Analysis of the number of prophage genomes annotated in the dataset. A. Linear regression analysis of the total number of prophage-like elements in function of the assembly size in bp. Phylogroups are color coded as follows: PG0 in teal, PG1 in violet, PG2 in orange, PG3 in blue, Pg4 in yellow, PG5 in light green, PG6 in pink, PG7 in purple, PG10 in olive, and PG13 in dark green. The correlation coefficient R^2^ was 0.099 indicating a very weak correlation between the two variables. The p-value (F-test) was <0.0001. B. Outlier boxplots of the total number of prophage-like elements per phylogroup. Phylogroup 6 encoded on average the most prophage-like elements (Tukey test with correction for multiple comparisons p-value < 0.05).

Incorporating the phylogroup in the model, we found that phylogroup 6 (Group A) and 4 (Group AB) carry on average the most genomic parasites, followed by phylogroups 2 (Group BC) and 3 (Tukey test, p-value < 0.05; Fig. 4B). Interestingly, phylogroup 1 contained on average the least number of genomic parasites in their genomes. As this phylogroup encoded the largest number of defense mechanisms, we hypothesized that the number of genomic parasites declines with an increasing number of phage defense mechanisms. However, we found very weak correlations between the two variables for all phylogroups (R2<0.15 and p-value>0.1), suggesting that the number of defense mechanisms is a poor predictor for the number of prophage-like elements (Supplementary Figure 6). Next, we assessed whether the number of prophage-like elements differed when a certain defense mechanism was encoded by an isolate. As such, we found that when an isolate encoded an Abi mechanism, the number of prophage-like elements increased (one-way ANOVA, p-value = 0.0402). Similarly, our analysis demonstrated that in case of encoded DRT, Druantia, DSR, Hishiman, Kiwa, PARIS, ParC, RloC, ShosTA, and SspBCDE mechanisms, the number of prophage-like elements were significantly higher compared to isolated without the respective mechanisms (one-way ANOVA, p-value < 0.05). On the other hand, when isolates encoded Gao, Menshen, Shango, or Wadjet systems, the number of prophage-like elements was significantly lower (one-way ANOVA, p-value < 0.05). Looking at the number of intact prophages, we observed similar trends with a significantly decreased number of prophages when an isolate encoded Abi, DarTG, Gao, Menshen, Olokun, Shango, Wadjet, and Zorya systems. The number of prophages was significantly higher in case of the presence of CRISPR-Cas, dCTP aminase, DRT, Druantia, Hashiman, Kiwa, Mokoshen, NRL, PARIS, ParC, RloC, and ShosTA mechanisms. Interestingly, Abi mechanisms popped up in our analysis as related to a decrease in the total number of intact prophages, and an increase in the total number of prophage-like elements.

Overall, there was no clear relationship between the prophage occurrence and *P. syringae* phylogroups. Since the phylogenomic model obtained for *P. syringae* was highly congruent to the phylogroups, the prophage frequency might be poorly linked to the bulk genetic background of *P. syringae*. In a few cases, a differential distribution of prophages within *P. syringae* was observed, such as in bacteria from phylogroup 10, which showed the absence of intact prophages, and some clades of phylogroups 1 and 2, which showed a higher frequency of such elements.

We employed a combination of vConTACT2 and VIRIDIC to access the sequence diversity of prophages. While vConTACT2 generates sequence clusters based on protein-sharing between phages (Figure 5), VIRIDIC uses local nucleotide alignments (BLASTn) to group sequences into species clusters and genus clusters according to user-defined thresholds of intergenomic similarities. The largest protein-sharing cluster generated by vConTACT2 contained most of the prophage-like elements, including nearly all intact sequences but also defective sequences, meaning that most prophage-like elements share some highly similar genes. Furthermore, this cluster grouped prophages harbored by *P. syringae* from all phylogroups. Three other clusters contained a reasonable number of prophage-like elements and were highly homogeneous regarding both sequence completeness and *P. syringae* phylogroup [top right, Fig. 5]. Visualizing only the intact prophages, we distinguished seven clusters. Remarkably, these clusters primarily contained prophages annotated within certain phylogroups. Yet, as illustrated in Figure 5, the different nodes were strongly interconnected, depicting a high degree of similarity between the different genomes. In total, VIRIDIC clustered the 2595 prophage-like sequences in 1282 species clusters, many containing only one sequence. The high sequence diversity was also evident by visualizing all pairwise comparisons between prophage-like elements, with few sequences sharing more than 65% of intergenomic similarities. Below this level, the prophage-like sequences are considered distantly related (24). Overall, the prophage-like elements occurring in a given species cluster had a narrow host range, occurring in a specific *Pseudomonas* subspecies. The exceptions included elements from ten species clusters, which were hosted by more than one *Pseudomonas* species.

**Figure 5.**
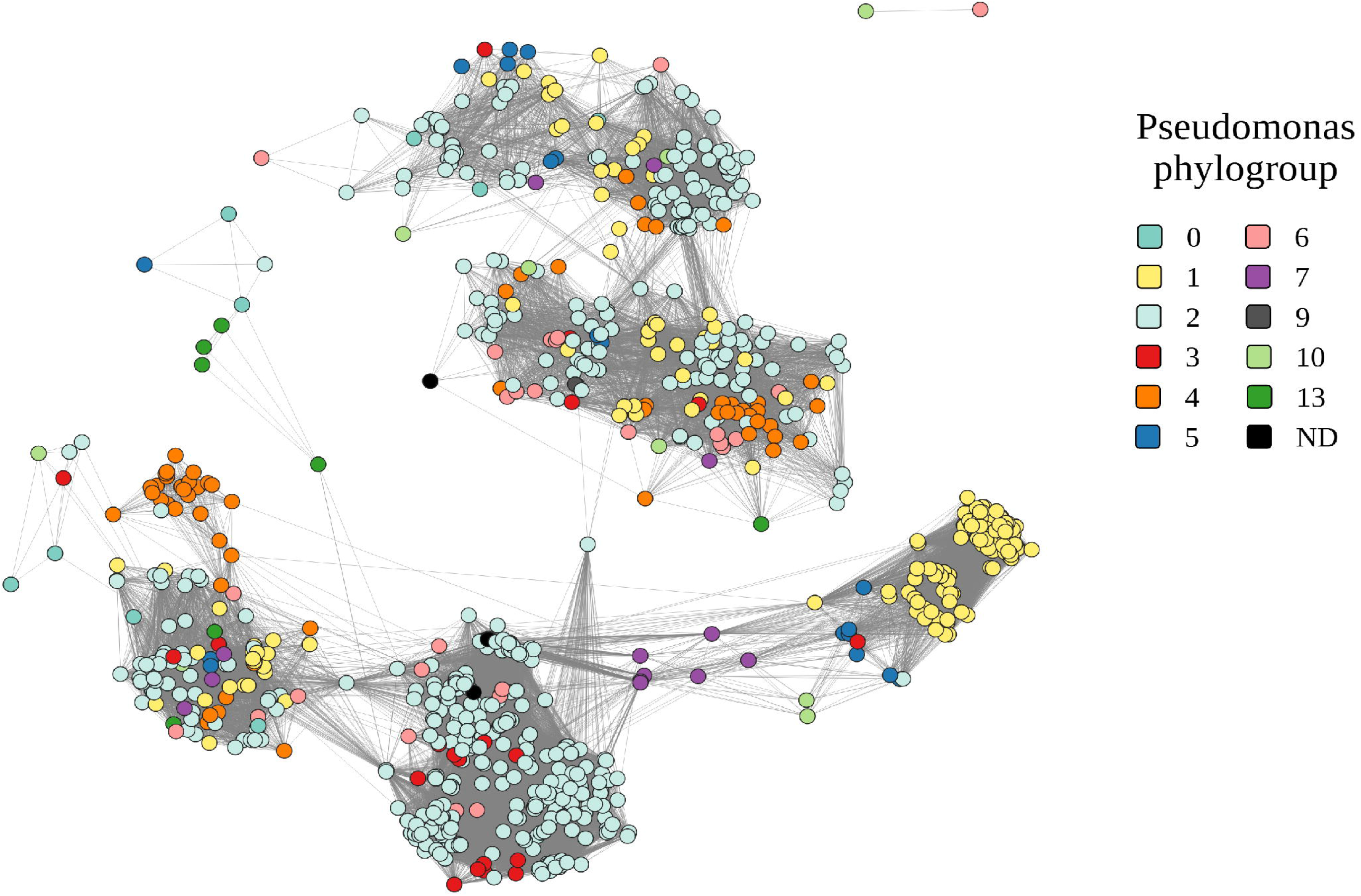
Gene sharing network between the different prophage genomes annotated in the *P. syringae* species complex. The nodes are colored along the phylogroups (PG0 teal, PG1 yellow, PG2 light blue, PG3 red, PG4 orange, PG5 blue, PG6 pink, PG7 purple, PG9 brown, PG10 light green, PG13 dark green and not determined in black). Every connection represents a similarity to another predicted viral sequence.

### The role of genomic parasites in the establishment of the *P. syringae* phage defensome and effectorome

Next, we mined all prophage-like elements for the presence of either phage defense mechanisms and/or effector genes. Based on this analysis, we found that 3.7% (254 defenses annotated in prophage-like sequences/7032 total defenses) and 1.4% (166 effectors annotated on prophage-like sequences/11748 total annotated effectors) of all phage defenses and effector genes are encoded by prophage-like elements as described by our analysis. Supplementary Figure 7 gives an overview of the relative abundance of the different phage defenses and effector proteins encoded by these genomic regions. To determine how potentially mobile these defenses and effectors are, we separated the prophage-like elements into cryptic prophages and viral satellites (which are typically not able to disperse on their own) and intact prophages that encode all the genetic information necessary to excise from the host genome and enter a lytic replication cycle. Based on our dataset, we observed that intact prophages also encode both phage defenses and effector proteins as illustrated by the red stars in Figure 3 and Figure 4. In total, we found that 10% of intact prophages encode either phage defenses and/or effector genes (7% encodes phage defenses 62/860 and 3% encodes effector genes 25/860).

Given the relatively low taxonomical distance among the *P. syringae* isolates within our dataset, we expected a taxonomical relationship among the prophages we identified. To this end, we compiled all the intact prophage genomes that encoded either phage defenses, effector genes, or both and performed a taxonomical analysis of the phages that carried them. Indeed, based on a Viptree and VIRIDIC analysis, we observed that the prophages clustered together into two main lineages, with one outlier (Supplementary Figure 8). One of the main lineages represented a novel group of phages that cluster within the *Peduoviridae* family. The second lineage was related to other, previously observed *P. syringae* phages, including *P. syringae* pv. *morsprunorum* phage MR15, *P. syringae* pv. *actinidiae* phages psageK9, PhiSA1, PhiAH14a and *P. syringae* pv. *tomato* phage Medea1. This latter clade split out into two subgroups. As summarized in Figure 6, the P2-like phages were mainly occurring in isolates belonging to phylogroup 2 (76% 25/33). In case of the lambdoid phages, one subgroup occurred mainly in phylogroup 2 (50%) followed by phylogroup 4 (27%). The other subgroup was primarily associated with phylogroup 4 (41%) and phylogroup 1 (37%).

**Figure 6.**
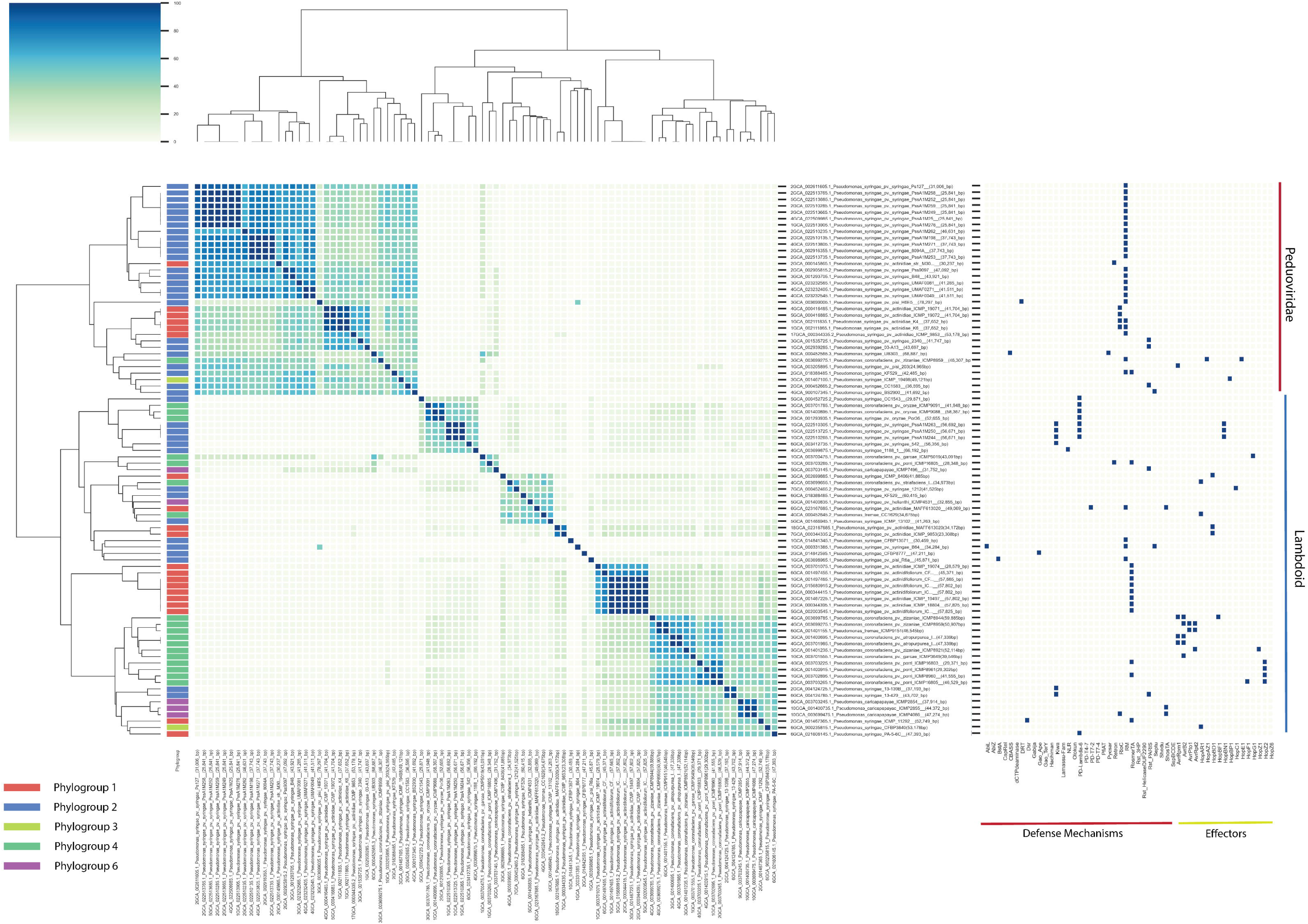
Taxonomical analysis of the prophage genomes as determined by a pairwise comparison of intergenomic similarity (VIRIDIC analysis – left panel) and absence/presence matrix of the different phage defense mechanisms and effector genes encoded by the prophage genomes (right panel). The different prophage genomes were clustered according to their genome similarity (100% similarity is blue, 0% similarity is light green). The phylogroups are color coded: PG1 red, PG2 blue, PG3 yellow, PG4 green, and PG6 purple. The prophages split out in two main clusters: the *Peduoviridae* and the lambdoid phages as determined by a VipTree analysis. Within the lambdoid phages, two clusters can be distinguished.

The most abundant phage defenses encoded within the prophage genomes were restriction modification systems and RosmerTA. While the former is most abundant in the *Peduoviridae*, the latter is most prominent in the second clade. Overall, we found significant differences between the number of phage defenses that were genomically- or prophage-encoded (Wilcoxon signed rank test; p-value < 0.001). This suggests that some mechanisms might be more likely to be carried by prophages than by chance. Indeed, we found that genes encoding RosmerTA, Kiwa, Rst_Paris, ShosTA, and PD-Lambda-6 were more abundant in prophage genomes than in the general genome, while Septu, Gabija, Retron, and CBASS mechanisms were enriched in the genome compared to prophage (Fig. 7A).

**Figure 7.**
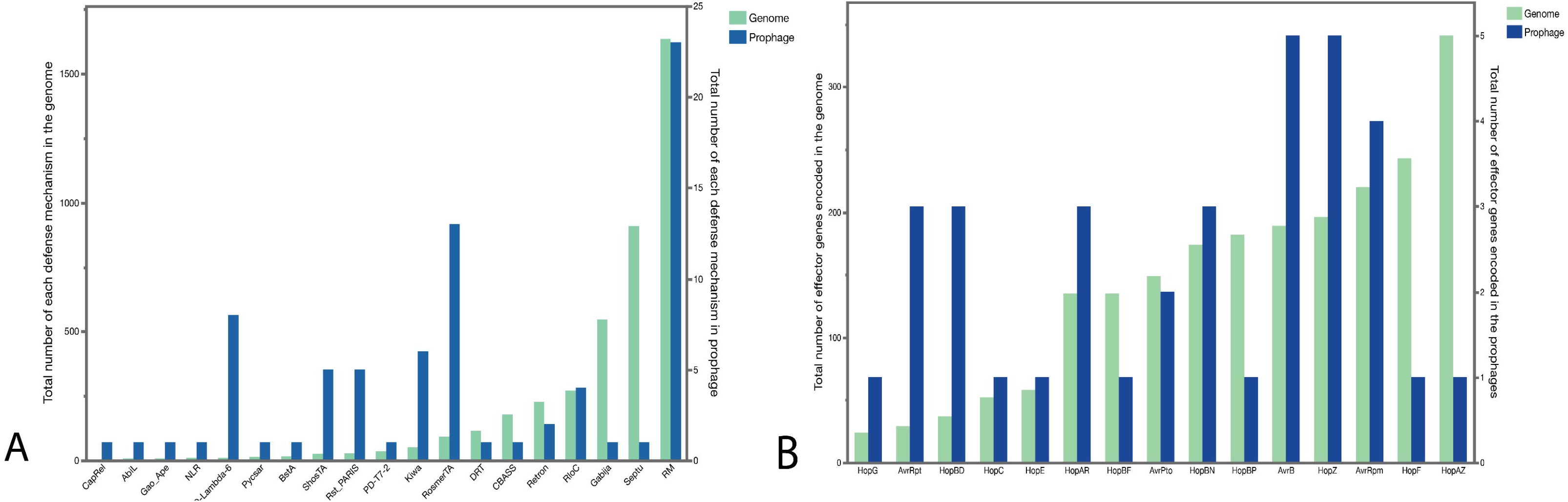
Total abundance of phage defenses (A) and effector genes (B) encoded by the prophage genomes. The left y-axes depict the abundance of the mechanisms in the genomes (teal) and the right y-axes graphs the total abundance in the prophage genomes (blue). Based on Wilcoxon signed rank tests, certain phage defense mechanisms and effector genes were enriched in the prophage genomes (p-value < 0.001).

Regarding effector genes in our dataset, we observed that lambdoid phages were the ones primarily carrying effector genes, with only a minority of the *Peduoviridae* annotated to contain effector genes (3/33). We found that *avrB*, *hopZ*, and *avrRpm* are relatively abundant within intact prophage genomes. Other effector genes that are common in *P. syringae*, like *hopAZ* and *hopF*, are less abundantly present in the prophage genomes. Again, based on a Wilcoxon signed rank test, we found a significant difference between the types of effectors observed in prophage compared to the genomic background of *P. syringae* (p-value <0.001). In particular, AvrB, HopZ, AvrRpm, AvrRpt, HopBD, and HopAR were more abundant in prophage genomes. This further suggests that there is an enrichment effect of specific genes in prophages that are likely not accumulated by chance and therefore potentially under positive selection to be mobile among strains (Fig, 7B).

## Discussion

Like all organisms on Earth, bacterial populations are constantly evolving in response to the abiotic and biotic environment around them. For bacterial pathogens, the biotic selection pressures are likely particularly intense, including those imposed by other microbial taxa, such as phages, and those imposed by the host organism’s immune responses. To cope with these - potentially conflicting - selection pressures, the mobilome offers a rapid and effective mechanism to both spread important traits within a community and increase the flexibility of bacterial evolution. We took advantage of the well-studied and highly sequenced *P. syringae* species complex to determine the importance of prophages in shaping bacterial responses to viral and host-associated selection pressures.

Looking at the whole *P. syringae* species complex and focusing on genes related to phage defenses and effector genes, we found that the isolates in this study harbor quite elaborate phage defensomes and effectoromes. While the pandefensome of the species complex consists of 139 different defense mechanisms, with an average of 12 defense mechanisms per genome, the average number of unique effector genes in *P. syringae* isolates clearly depended on whether isolates were host-associated (average is 12 defense mechanisms) or environmental isolates (average is 7 defense mechanisms; Figure 1), slightly higher than previous estimates from across a highly diverse panel of 21,364 bacterial genomes (34). For other plant-associated bacteria specifically, this number has been shown to be as high as 10 mechanisms for some *Ralstonia* and *Xanthomonas* isolates (34). In sharp contrast to other bacterial species (35, 36), and even other Pseudomonads, CRISPR-Cas does not appear to be an important phage defense strategy in the *P. syringae* species complex (37, 38). Instead, we demonstrated that *P. syringae* relies on other defense mechanisms, such as restriction enzymes and abortive infection strategies, to cope with phage predation.

Despite there being a clear phylogenetic signal for the absence versus presence of phage defense mechanisms encoded by the *P. syringae* species complex, we observe a clear role of horizontal gene transfer in the establishment of the phage defensome, similar to other bacterial taxa (Figure 3 and Figure 4) (39). As prophages are an integral part of the mobilome, we assessed the potential importance of prophages in carrying, and potentially moving, these defense and effector genes among bacteria and we next quantified the number of prophages carried within the *P. syringae* species complex. We find that there were, on average, 4 prophage-like elements encoded per genome across the 590 isolates of *P. syringae,* demonstrating the abundance of prophage-like elements in the genomes of these isolates (Figure 5). This average number is comparable to other phytopathogenic bacteria such as *Erwinia amylovora,* which also carries a panel of 4 prophage-like elements per genome (36). We further observed that the average number of prophage-like elements, phage defenses and effectors encoded within both the genome and the mobilome differs across the phylogroups. Interestingly, the isolates with the highest number of prophage-like elements, phage defenses and effectors per genome are host-associated, similar to encoded viral elements in the genome (Figure 1 and Figure 5). Hence, it seems that in the host environment, bacteria are more exposed to phages in general, contributing to the observation that these isolates contain both higher numbers of prophages and host defenses. Of all the phylogroups, phylogroup 1 (primarily consisting of *P. syringae* pv. *actinidiae* infecting kiwi vines), was observed to have the most elaborate phage defensomes and effectoromes.

Because prophages have the capability of killing their host cells, the widespread maintenance of these genomic parasites suggests that they likely also offer a selective advantage to their host cells. To estimate this, we specifically focus on genes that play a role in bacterial interactions with lytic phages (phage defense systems) and those involved in virulence on plant hosts (the effectors). Within our analysis, we observed a total of 83 unique prophages encoding a phage defense mechanism and/or an effector gene, and hence potentially influencing their host’s interaction with the other phages and/or the plant host. The proportion of prophages harboring either phage defense mechanisms or effector genes is quite consistent across PG1, PG2 and PG3 with 13% (21/160), 13% (38/298), and 14% (2/14), respectively. PG6 and PG4, on the other hand, have drastically more prophages encoding these factors with 30% (3/10) and 65.5% (19/29) of prophages carrying phage defense mechanisms and/or effector genes, respectively.

Looking more closely at the individual effector genes and defense mechanisms encoded by prophages, we found a clear enrichment effect of effector genes and phage defense mechanisms in prophage genomes that is likely not the result of chance. This finding suggests that also in case of *P. syringae,* lysogenic conversion may contribute to the overall virulence of the species. Strikingly, this effect seems quite apparent in phylogroup 4 in which there were 19 different prophages annotated to carry at least one effector gene (and/or defense mechanism; Figure 7). Based on this observation, we can hypothesize that the virulence of isolates from phylogroup 4 is quite dependent on prophage-encoded effector genes. Indeed, a comprehensive study of the association between prophages and effector genes in *P. syringae* pv. *morsprunorum* isolates belonging to phylogroup 4 demonstrated the successful distribution of a prophage encoding HopAR1 from epiphytes within the phyllosphere community to pathogenic *P. syringae* pv. *morsprunorum* isolates (33). Our taxonomical analysis of the prophage genomes further indicated that the prophages encoding effector genes were primarily lambdoid phages. The role of this taxonomic group of phages in enhancing the virulence of their host was previously described in *E. coli* were these phages were found to encode the Shiga Toxin (Stx) and hence contribute strongly to the pathogenicity of the host (40). Our results emphasize the overall role of lambdoid phages in driving pathogen-host interactions in another bacterial species, raising the question of the overall relevance of these phages in pathogenesis of pathogens and pathogen evolution in general. Alongside the effector genes, many antiphage mechanisms have been found to be encoded in the prophage genomes that defend the host (18, 20, 41–44). We find that the *Peduoviridae,* in particular, play a large role in encoding phage defenses, including by carrying restriction enzymes and other phage defenses that underlie superinfection exclusion (Figure 7 and Figure 8). Similarly, in *E. coli* P2-like phages, together with P4-like satellites, carry hotspots encoding diverse anti-phage systems, suggesting the importance of these phages in the establishment of the host’s phage defensome (45). Hence, we can hypothesize that this family of phages likely drives the evolution of their hosts and interactions within the microbial community, in particular the sensitivity of isolates to other phages within the viral community. Since these phages seem quite diversely spread among isolates within phylogroup 2, we can further hypothesize that there is an intricate relationship between the phages and isolates from this specific phylogroup.

Altogether, our results, emphasize the role of specific prophages on the evolution of their host. However, we observe that not all families of prophages contribute equally to provide their host with an evolutionary advantage towards phage predation and virulence. Lambdoid phages and members of the *Peduoviridae* appear to influence the virulence of their hosts across bacterial species, allowing us to hypothesize on the intimate relationship between these viruses and their hosts. Our analysis, however, strongly depends on the quality of the genomes and the correct annotation of effector genes, defense mechanisms, and prophage genomes. With upcoming advances in both long read sequencing and the accuracy at which prophages are annotated by recent tools such as geNomad (46), we predict that the accuracy of analyses like this one will improve significantly in the foreseeable future. Furthermore, as this study employs a comparative genomics approach, we have no insights on the actual transcription of the defense mechanisms, nor the effector proteins. Future endeavors should focus on the *in situ* expression of these genes to unravel how they steer the evolution of and interaction between the phages, bacteria, and ultimately the higher organisms. Similarly, future research should focus on the activity of the prophages described in this analysis and their ability to distribute genes from one host to the next. As mentioned earlier, Hulin and colleagues have presented the first endeavors to demonstrate the distribution of virulence genes between *P. syringae* isolates by prophages (33). Yet, knowledge on the expansion of genes that are distributed by prophages among other genomic parasites within microbial communities is lacking. As such, our analysis, through comparative genomics, provides one of the first critical steps to understand which genes are distributed and ultimately the role of genomic parasites in the evolutionary trajectory of their host.

## Conclusion

Ass genomic parasites of bacteria, prophages are an important part of the bacterial mobilome, but can also kill their host cells under environmental stress. The fact that these Trojan horses are widespread in bacterial genomes implies an evolutionary advantage for maintaining these parasites in the genome. In our study, we demonstrated the role of specific viral taxa in driving the effectorome and phage defensomes in *P. syringae*, critical in steering bacterium-plant and bacterium-virus interactions. We further provide statistical evidence that host-associated isolates have more elaborate phage defensomes, effectoromes, and prophage-like elements incorporated in their genomes. This raises questions on the role of these viruses in the dispersal of these genes within the broader host-associated microbial community as well as the frequency of temperate phage infections within the microbiome and how these viruses are shaping the evolutionary trajectory of pathogens and other members within the microbial community. Our study also provides insights into the role of phage defenses, characterized by an incredible diversity and distribution in bacterial genomes, in phage-phage competition alongside bacterial defense against phage infection.

## Materials and Methods

### Data acquisition, quality control and phylogenetic analysis

A dataset of 630 draft genomes of the *Pseudomonas syringae* species complex was formed by extracting all strains available (in January 2023) from the NCBI taxonomy database. The dataset was curated for doubles and the quality of each strain’s genome was assessed using BUSCO (v5.4.7), and strains with a score of at least 0.95 were used in all further analyses (47). A phylogenetic analysis on the soft-core genome of the different isolates was performed by means of PPanGGOLin (v1.2.105) (48).

### Annotation of bacterial genomes and analysis of the phage defensome and effectorome

The draft genomes were annotated with Prokka (v.1.14.5) for an open reading frame prediction (49). The FASTA Amino Acids files were mined for phage defense mechanisms by means of DefenseFinder (v.1.0.9) (34) with default parameters. The output files were compiled, and different defense mechanisms were summarized per draft genome. Similarly, the genome faa files were mined for the presence of effector genes by means of phmmer (e-value < 1E-60) using the database as described by Dillon *et al.* (29). Statistical analyses and data visualization were performed by means of the JMP Pro16 software. An absence/presence matrix was constructed, and a hierarchical clustering was performed by means of JMP Pro16. The similarity of defensomes between phylogroups was evaluated by an analysis of similarity (ANOSIM) using the VEGAN package in R (50).

### *In silico* analysis of horizontal gene transfer

A RecPD (v0.0.0.9000) analysis was used to calculate the degree at which every defense mechanism was transferred through horizontal gene transfer in the *Pseudomonas syringae* species complex (51). The phylogenetic tree of the *Pseudomonas syringae* species complex and the absence presence matrix generated from DefenseFinder were provided to the RecPD program as input. The nRecPD statistic for each defense mechanism was calculated by dividing the total branch length in which the defense mechanism was transferred vertically by the total length of branches of the tree rooted at the defense mechanism’s most recent common ancestor (34, 51). All defense mechanisms with a nRecPD score lower than 0.25 were used in all subsequent analyses of horizontal gene transfer within the *Pseudomonas syringae* species complex.

To this end, the genomic islands as determined by the PPanGGOLin (v1.2.105) analysis were mined for the defense mechanisms with the highest probability of being dispersed by horizontal gene transfer. A Blast database containing a random instance of every defense mechanism with a nRecPD score lower than 0.25 was constructed using the makeblastdb application (v2.14.0) with default parameters for nucleotide sequences. All DNA sequences of each defense mechanism found in the Pseudomonas syringae species complex were obtained through a Blastn analysis of these genomic islands using this database of defense mechanisms, an e-value of 1E-6, and default values for all other parameters.

Using MegaX (v11.0.11) (52), a phylogenetic analysis was performed to validate the nRecPD statistic for measuring horizontal gene transfer. For each defense mechanism, a multisequence alignment was performed at the protein level using the MUSCLE algorithm in MegaX with default parameters on every instance of this defense mechanism found in the pangenome. A phylogenetic tree was built from every multisequence alignment using maximum likelihood estimation in MegaX with default parameters. The phylogroup of every strain was annotated onto each phylogenetic tree using the ggtree R package (53).

### Prophage analysis

The *Pseudomonas* genome sequences were submitted to PHASTER (54) via the API to detect prophage-like sequences. Since many *Pseudomonas* genome assemblies were fragmented in multiple separated fasta sequences, we analyzed if the questionable and incomplete prophages identified by PHASTER were near the contig edges. In this scenario, the prophage-like sequences could be artificially fragmented, hindering the completeness estimated by PHASTER. Following this, the putative questionable or incomplete prophages occurring in 1000 bp or less from contig edges and sizes less than 10 kb were considered a possible artificially fragmented sequences.

All identified prophage-like sequences were submitted to Prokka v.1.14.5 (49) for open-reading frame prediction (parameters: --kingdom Viruses --gcode 11). The heatmaps showing prophage occurrence were plotted within a *Pseudomonas* soft core genes phylogenomics. The plot was generated using the “ggtree” package (53) in the R environment (R Core Team, 2013).

The gene-sharing networks of the prophage-like sequences were conducted using vConTACT2 (55)(parameters: --rel-mode ‘Diamond’ --db ‘None’ --pcs-mode MCL --vcs-mode ClusterONE). Cytoscape v.3.8.2 was used for networking visualization. Node colors and shapes were set based on *Pseudomonas* phylogroups and prophage sequence completeness, respectively.

We used VIRIDIC (56) to obtain an overview of whole-sequence similarity between prophages-like sequences. VIRIDIC uses local alignments (BLASTn) to calculate nucleotide-based intergenomic similarities and sequence clustering in “species_cluster”, as long as they share sufficient intergenomic similarities (default threshold of 95% for species clustering). The plots were generated using the “circlize” package (57) in the R environment.

Similarly, the prophages that were annotated to encode effector genes and phage defense mechanisms were taxonomically classified by means of VipTree (58) and the intergenomic similarity was calculated with VIRIDIC and visualized with the cluster heatmap package from Seaborn (v0.13.0) (59).

## Acknowledgements

This research was funded by an NSF and USDA/NIFA Career Award (NSF # 1942881, USDA/NIFA # 1024053). We further acknowledge the Coordenação de Aperfeiçoamento Pessoal de Nível Superior-Brasil (CAPES) PrInt #88887.697048/2022-00, Fundação de Amparo à Pesquisa do Estado de Minas Gerais (FAPEMIG) Fapemig #APQ-00661-18, INCT Plant Pest Interactions #406440/2022-0 and Conselho Nacional de Desenvolvimento Científico e Tecnológico (CNPq #313528/2021-7). BK is a Chan-Zuckerberg San Francisco Biohub investigator.

