## Supplementary figures for "Evidence for the key roles of the *Pseudomonas syringae* mobilome in shaping biotic interactions" (from another version of this preprint: version 1, 2024-03-20)

Supplementary information Supplementary Figures

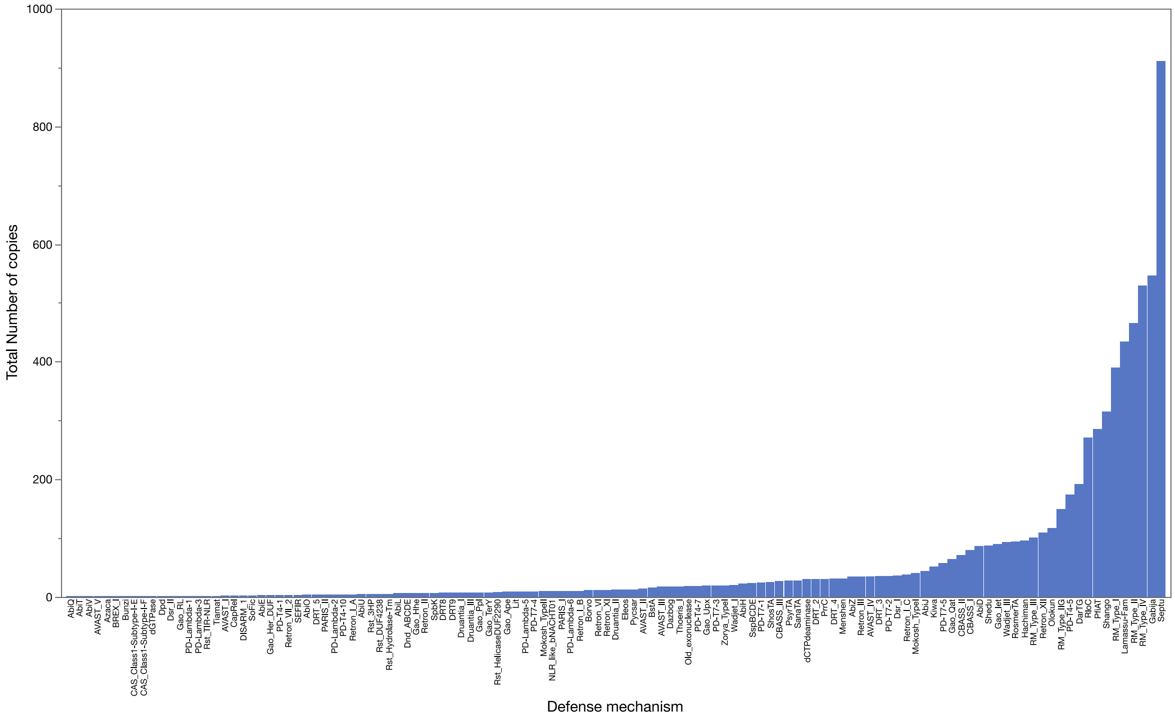


Supplementary Figure 1 Overview of the full phage defensome of the *P. syringae* species complex with septu and restriction enzymes the most abundant phage defenses.

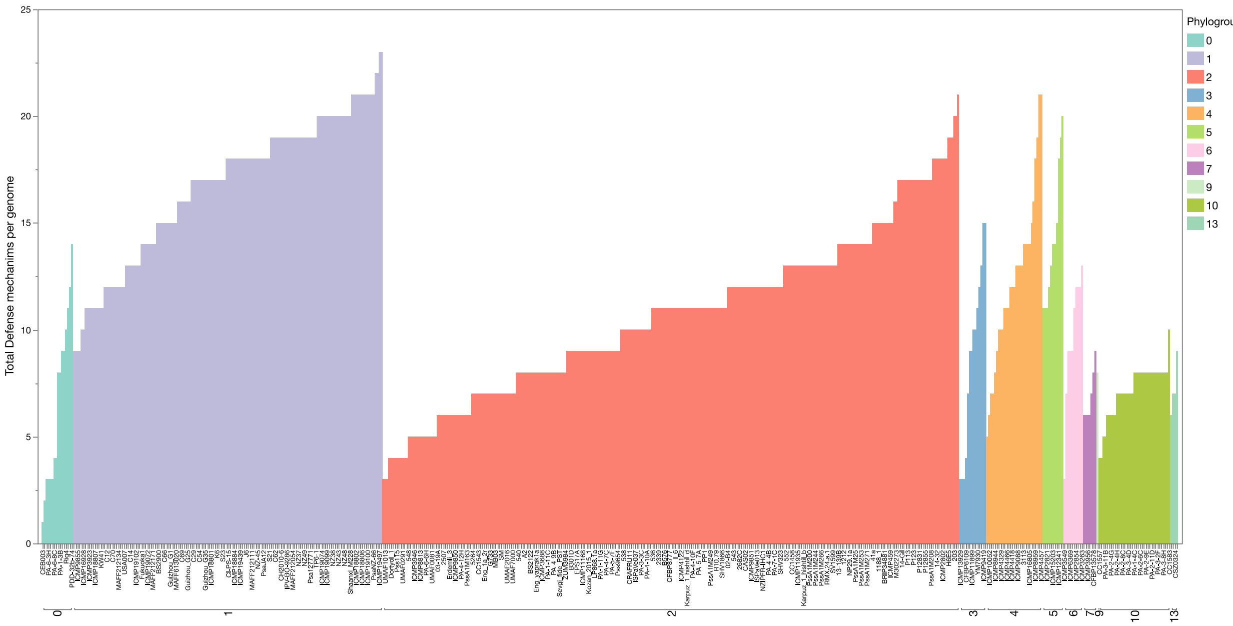


Supplementary Figure 2 Overview of the number of defenses encoded by the different isolates organized per phylogroup. In cyan PG0, purple PG1, orange PG2, blue PG3, yellow PG4, light green PG5, pink PG6, violet PG7, green PG10, and teal PG13.

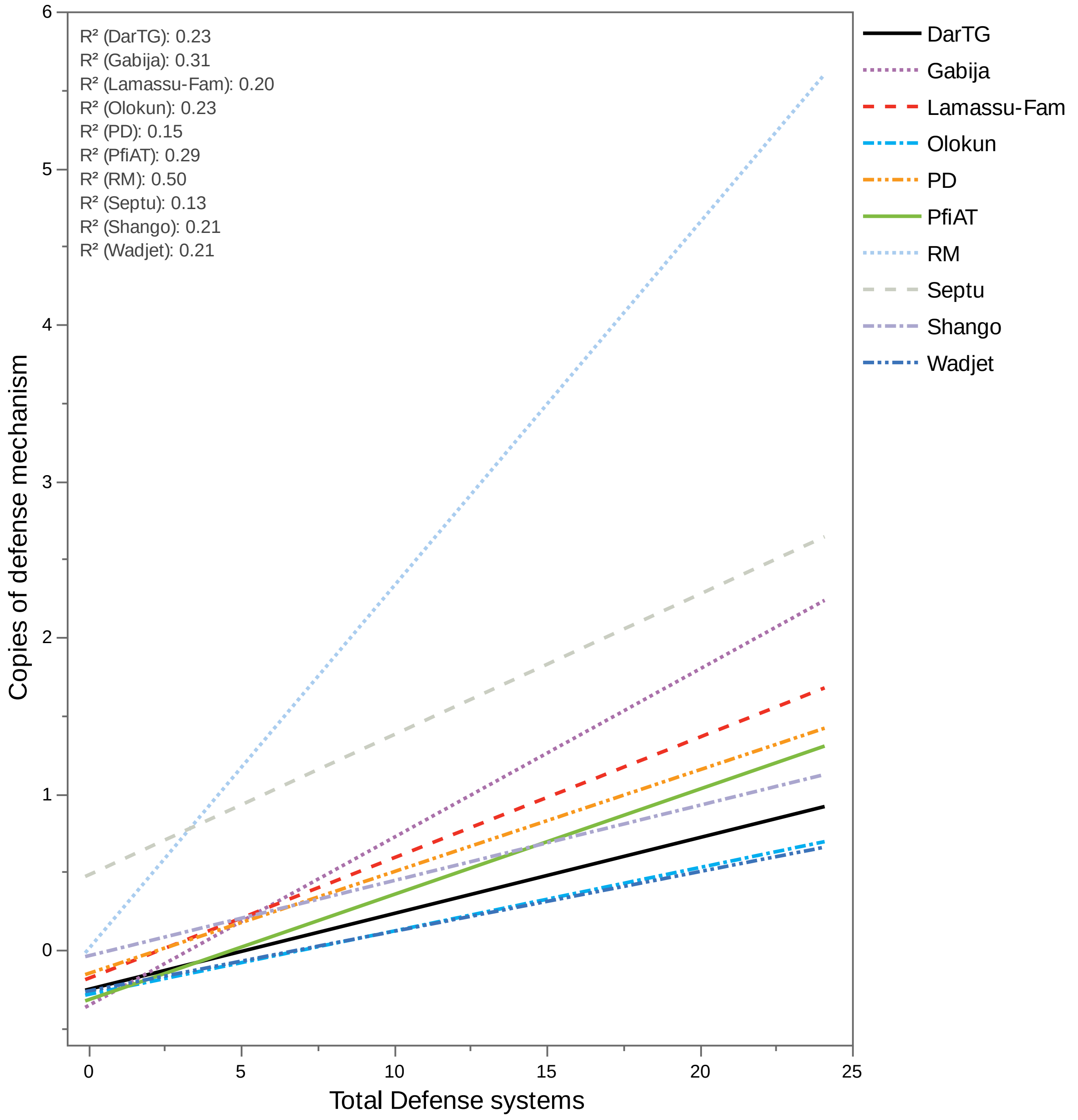


Supplementary Figure 3 Linear regression of the copy number of a specific defense mechanisms in function of the total number of defense systems encoded by a specific isolate. The R2 values are indicated in the upper right corner of the graph. In dark blue Wadjet, purple Gabija, red Lamassu-Fam, blue Olokun, oranje PD, green PfiAT, light blue RM, grey Septu, and gris violet Shango.

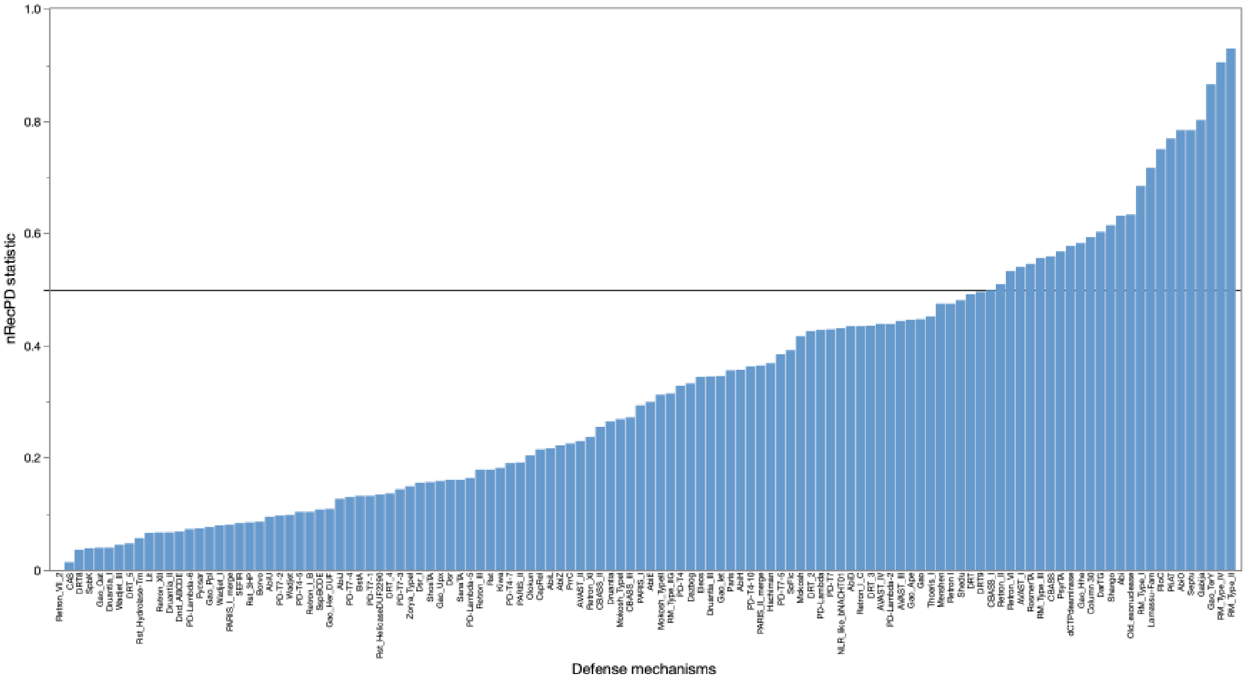


Supplementary Figure 4 nRecPD values for the different phage defense mechanism calculated based on the phylogenetic distance as determined by a core genome alignment of the different isolates. The threshold of 0.5 indicates mechanisms that are more likely to be distributed through horizontal gene transfer (value closer to 0) or vertically inherited (value closer to 1).

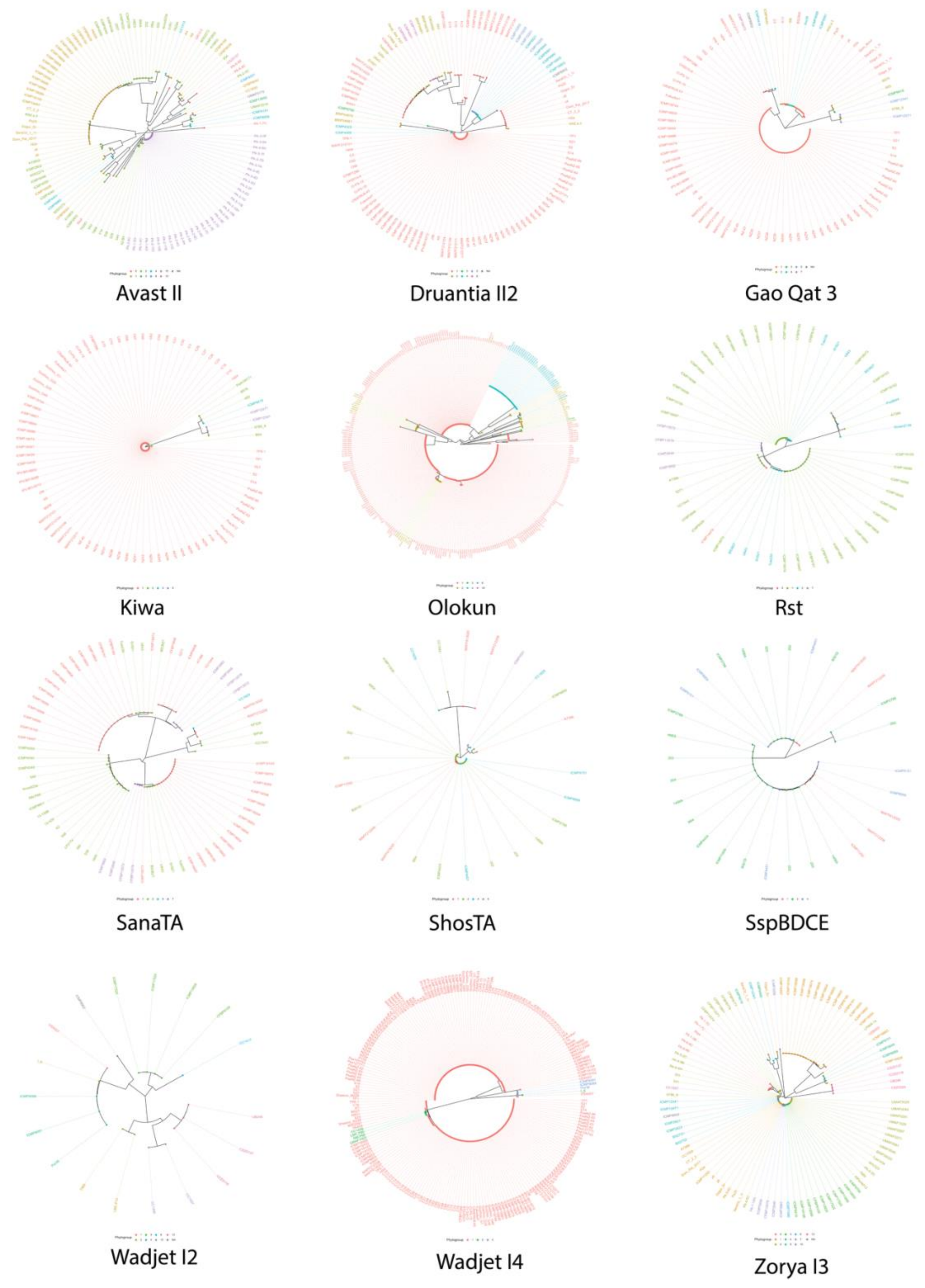


Supplementary Figure 5 Maximum likelihood trees midpoint rooted (with 1000 bootstraps) for the concatenated amino acid sequence of the full phage defense mechanism. The different phylogroups were color coded as indicated below every tree.

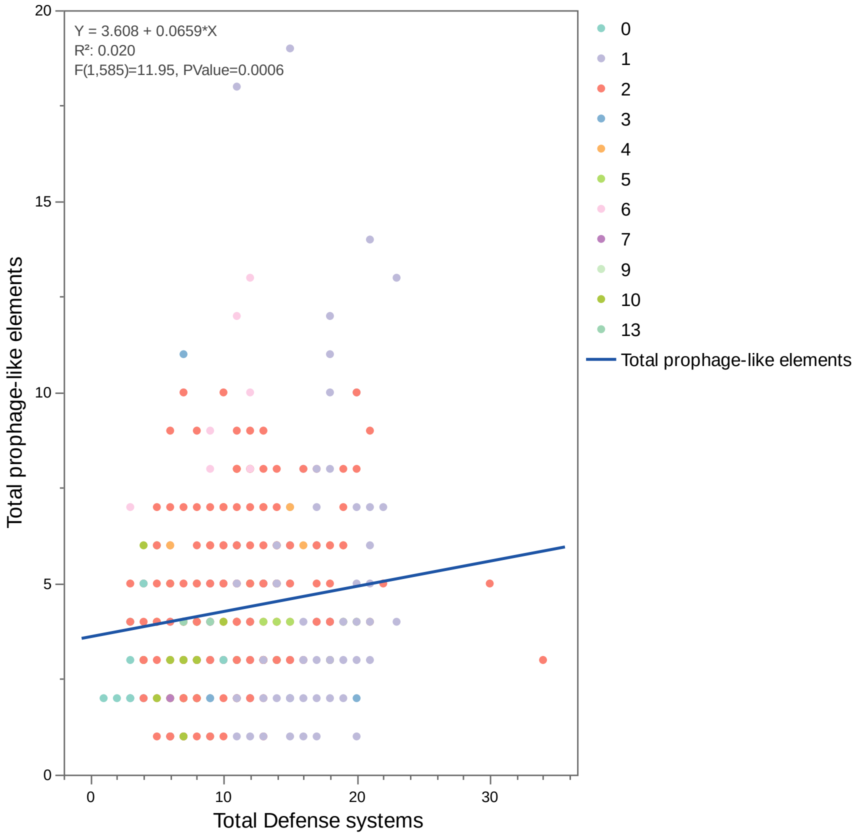


Supplementary Figure 6 Linear regression of the total number of prophage-like elements encoded in function of the total number of phage defenses per isolate. A rather low $R^2$ value indicates a weak correlation between the two variables, suggesting that the total number of phage defenses is a poor predictor for the number of genomic parasites in the genome.

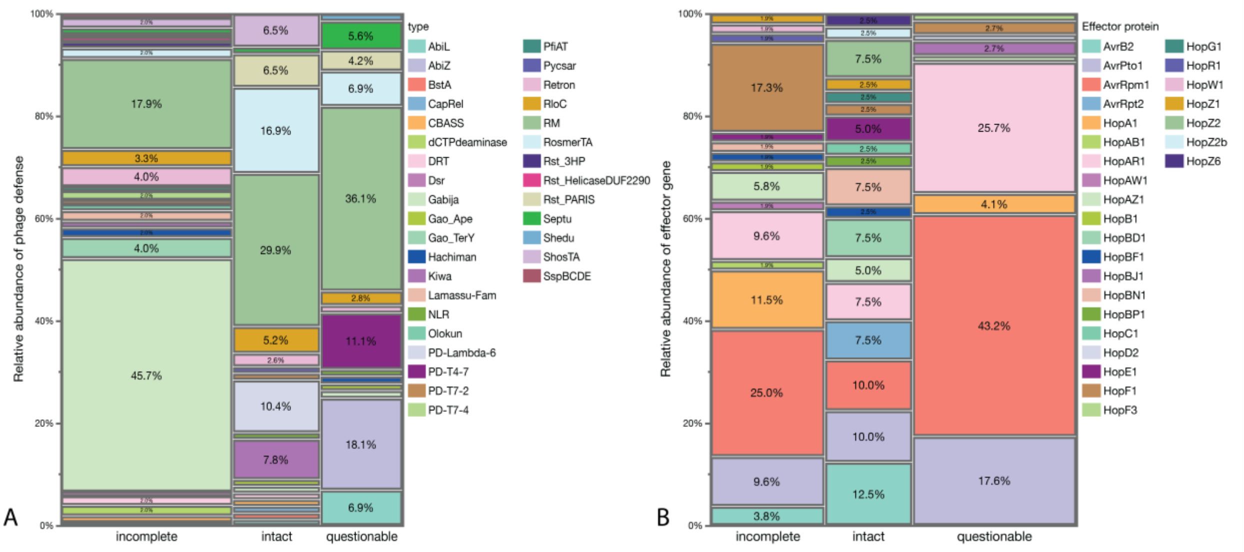


Supplementary Figure 7 Relative abundance of the different defense mechanisms (panel A) and effector genes (panel B) encoded on the prophage-like genetic elements organized in three groups (incomplete prophages, intact prophages, and questionable prophages).

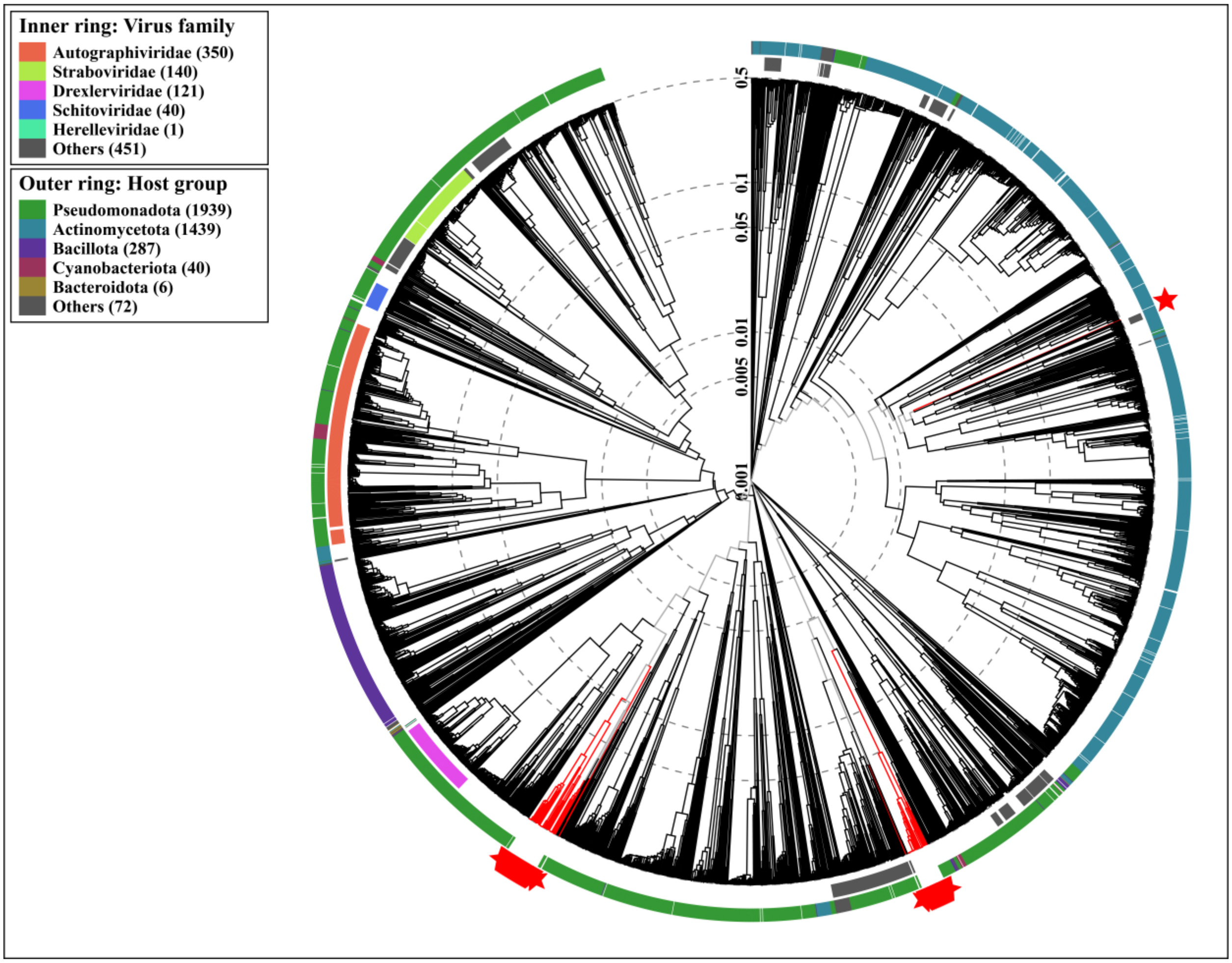


Supplementary Figure 8 VipTree analysis of the prophage genomes encoding effector and/or phage defenses. Genomes from this analysis are indicated by the red stars. Two different clades can be distinguished and one orphan phage.
